# Insulin regulates systemic lipid traffic in starving animals

**DOI:** 10.1101/2022.05.09.491148

**Authors:** Laura Christin Trautenberg, Maria Fedorova, Marko Brankatschk

## Abstract

In periods of malnutrition starving organisms initiate the reorganization and degradation of cellular lipids. Yet, the build-up of lipid stores is reported in animals entering early starvation stages. Anabolic lipid metabolism is regulated by many circuits including the insulin/insulin-like growth factor signaling (IIS) cascade. The regulative role of IIS ligands in food deprived animals is often neglected, since supporting evidence for this notion is largely missing.

Here we show that in starving adult Drosophila melanogaster IIS is required to stabilize the transport of circulating lipids and sugars, and is essential for the initial increase of lipid stores in fat body cells. Further, we demonstrate that the *Drosophila* insulin-like peptides (dIlp) 2 and 5 regulate the cellular IIS cascade differentially. To its end we speculate that each hormone initiates complementary measures that increase the starvation resistance of adult flies.

## Introduction

Metabolic homeostasis and adaption are vital in challenging environments. Insects, such as *Drosophila melanogaster*, consistently face nutritional scarcity because food sources are patchy in nature. One challenge is the maintenance of metabolic rates in order to withstand starvation periods in between intermitted foraging of adult flies.

The response to starvation can be defined as a series of metabolic switches required to fill the energy needs of organisms. During starvation low blood glucose levels are restored by the degradation of glycogen stores. Once glycogen is depleted, the metabolism transitions towards fat mobilization and oxidation to ensure energy supply across tissues. For the transition into fat metabolism the organism undergoes several processes. Initially, starvation is characterised by a switch from phospholipid to triacylglycerides (TAGs) synthesis for building energy stores in preparation of nutrient deprivation (Stout et al., 1976). The build-up of TAGs is considered a part of anabolic turnover and the regulating signaling cascades remain elusive. In prolonged starvation, lipolysis is used to degrade TAGs into diacylglycerols (DAGs) and fatty acids (FAs) (Lehmann, 2018). Dedicated transport systems are capable to traffic these TAG products to their destined cellular and tissue targets (Canavoso et al., 2001; Palm et al., 2012). For cellular traffic of lipids in intestinal or fat body cells, the microsomal triglyceride transfer protein (MTP) is required to load DAGs onto lipophorin (LPP) lipoprotein particles (Palm et al., 2012; Sellers et al., 2003). LPP particles are subsequently secreted into circulation delivering lipids to peripheral tissues. To produce energy from lipids, FAs become accessible to mitochondria by the activity of the carnitine palmitoyltransferase 1 and 2 (CPT1 and 2). Mitochondria degrade FAs in a process named beta oxidation resulting in the production of molecules suitable for energy production or precursors usable in the gluconeogenic pathway (Lehmann, 2018).

Both genes for MTP and CPT are targets of the forkhead transcription factor (FOXO). FOXO is the terminal transcription factor of the cellular insulin signal cascade (Puig et al., 2003). Explained in brief, in feeding animals the ligand bound insulin receptor activates the phosphoinositide-3-kinas (PI3K). PI3K converts the lipid phosphatidylinositol(4,5)bisphosphate (PIP2) into phosphatidylinositol(3,4,5)trisphosphate (PIP3) at the inner leaflet of the cellular plasma membrane and the latter lipid recruits the enzyme Akt (protein kinase B/PKB) (Scanga et al., 2000). Membrane bound Akt can be activated by the mammalian target of rapamycin complex 2 (mTORC2) by phosphorylation at the amino acid position Akt^SER505^ (Sarbassov et al., 2005), and in parallel targeted by the enzyme PDK1 at the Akt^THR342^ (Cho et al., 2001). Flies express three isoforms of the Akt protein, dAkt^66^, dAkt^85^ and dAkt^120^ (Andjelkovic et al., 1995). They are thought to be homologous to the three encoded genes in mice and humans, Akt1, 2 and 3 (Gonzalez and McGraw, 2009). Phosphorylated and therefore activated Akt phosphorylates FOXO repressing its transcriptional activity. Thus, hepatic MTP and CPT levels in fed animals are lower than in starving controls (Kamagate et al., 2008; Park et al., 1995).

Insulin/insulin-like growth factor signaling (IIS) is an evolutionary old circuit that is highly conserved throughout the animal kingdom. In *Drosophila*, IIS ligands are represented by several *Drosophila* insulin-like peptides (dIlps) that target one single receptor. In contrast, vertebrates produce one insulin peptide and multiple insulin-like growth factors (IGFs). Insulin shows the highest affinity to its receptor but is able to bind IGF receptors as well (Soos et al., 1993). In *Drosophila*, dIlps regulate shared and specific functions (Grönke et al., 2010). Given the diversity of these functions it is poorly understood how signal specificity can be produced with one common receptor. For instance, it was shown that DILP2 and DILP5 bind to the insulin binding protein ImpL2 (Honegger et al., 2008; Sajid et al., 2011), and it is likely that both DILPs share the binding site on the insulin receptor. Both *dIlps* are expressed in adult brain median neurosecretory cells, named insulin producing cells (IPCs). dIlp2 is viewed to signal as an equivalent to mammalian insulin. It affects adult lifespan and plays a role in sugar homeostasis (Grönke et al., 2010; Post et al., 2018). dIlp5 was suggested to regulate protein and lipid metabolism, but evidence remains correlative at best (Grönke et al., 2010; Semaniuk et al., 2018). However, in starving animals dIlp5 mRNA levels are strongly reduced, whereas dIlp2 mRNA yields remain unaffected (Bai et al., 2012; Ikeya et al., 2002; Sudhakar et al., 2020).

It was shown that overnight fasting increases the mRNA of *dIlp6* in the fat body (Bai et al., 2012). dIlp6 is structurally a rather distant insulin like peptide resembling more IGF than insulin (Grönke et al., 2010; Okamoto et al., 2009), and in starving animals dIlp6 improves the starvation resistance presumably by facilitating the FA degradation in oenocytes (Chatterjee et al., 2014). The *Drosophila* fat body functionally represents the white adipose tissue and liver of vertebrates. Fat body cells serve as glycogen and lipid storage and show endocrine activity required to adapt to metabolic changes (Arrese and Soulages, 2010). Storage lipids are neutral lipids deposited in lipid droplets which form prominent organelles in fat body cells. This gives the fat body a key role in regulating metabolism during nutritional hardships.

Nevertheless, canonical IIS is not considered essential in mammals. Evidence for this rests primarily on two findings: starving animals can develop insulin resistance and the production/secretion of insulin is downregulate (Fery et al., 1990; Gannon et al., 1996). Here we show Drosophila, that Δ*dIlp2* or Δ*dIlp5* are starvation sensitive compared to Δ*dilp3* or wild type controls. Both starving mutants show different circulating sugar and lipid yields indicating distinct roles for dIlp2 and dIlp5 in the mobilization of resources. Focussing on the fat body, we found that dIlp2 and dIlp5 are responsible for the degradation rate of TAGs stored in lipid droplets. However, we found that dIlp2 signalling is required to secrete sufficient amounts of lipoprotein particles into the hemolymph. On the other hand, dIlp5 is required to stabilize blood sugar levels. Moreover, we provide evidence that dIlp5 is signalling through phosphorylation of AKT^66^. We propose that dIlp2 and dIlp5 target the insulin receptor of fat body cells, but facilitate different signalling pathways downstream of the kinase AKT.

## Results

### dIlp2 and dIlp5 are required in starvation

DILP signaling is required to build fat stores. We speculated that the loss of dIlp2, 3 or dIlp5 in starving animals should be reflected in the survival of adults. To test the idea, we kept staged flies on agar plates and tracked their starvation resistance over time. We found that Δ*dIlp2*, Δ*dIlp2-3* and Δ*dIlp5* were more starvation sensitive than Δ*dIlp3* or controls (Figure 1A). It was shown that the ablation of insulin producing neurons results in developmental delay and growth retardation (Rulifson et al., 2002) and therefore, *dIlp* mutants could show starvation-related phenotypes due to low feeding activity. To assess the feeding frequency of Δ*dIlp2, 3* or Δ*dIlp5*, we recorded the foraging behaviour of all three mutants. We found that Δ*dIlp5* fed more often than Δ*dIlp2, 3* or controls (Figure 1B) indicating that starvation resistance differences in these mutants are not based on reduced food uptake.

**Figure 1.**
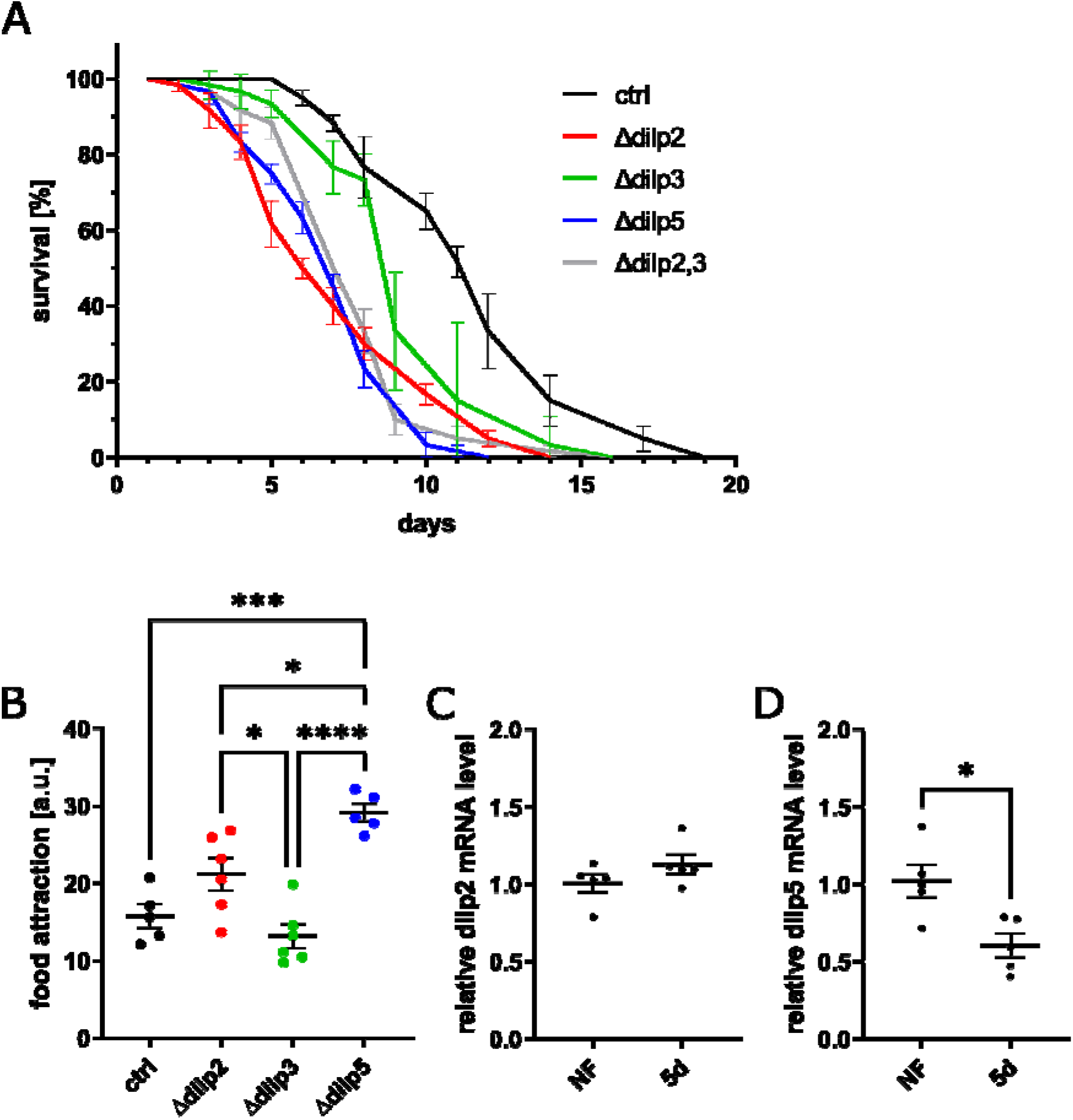
Starvation resistance of *dIlp* mutants is independent of preceding feeding behaviour. (A) Plotted is the survival curve of flies exposed to starvation on agar plates (n = 5). (B) Shown is the feeding behavior of *dIlp* mutants represented by video recorded attraction of flies to a normal food source. Shown is the mean±SEM, one-way ANOVA followed by Tukey’s multiple comparisons test (* p < 0.05, *** p < 0.001, **** p < 0.0001). (C, D) Shown is the relative mRNA level of dIlp2 (C) and dIlp5 (D) in fly heads of flies reared on normal food (NF) or starved for five days (5d) on agar plates. Shown is the mean±SEM, unpaired Student’s t-test (* p < 0.05).

It is reported that dIlp5 but not dIlp2 mRNA expression is downregulated in short-term fasting experiments (Bai et al., 2012; Ikeya et al., 2002; Sudhakar et al., 2020). To test, if under prolonged starvation dIlp2 and dIlp5 expression is changed, we decided to assess mRNA yields by qPCR. We found that in our assays the starving wild type animals showed reduced dIlp5 expression after 5 days, whereas dIlp2 mRNA yields remained unchanged with respect to fed controls (Figure 1C, D). Thus, in starving wild type animals relative higher DILP2 levels could possibly outcompete DILP5 at their targets.

Taken together, we confirmed that DILP2 and DILP5 regulate feeding behaviour and the prolonged starvation resistance of adult *Drosophila*. dIlp3 plays a minor role, since Δ*dIlp3* flies are more starvation resistant and *dIlp2-3* double mutants are not more starvation sensitive than Δ*dIlp2* or Δ*dIlp5* flies. However, starvation induced expression changes of DILP2 and DILP5 could indicate starvation-specific functional roles for either hormone.

### dIlp2 and dIlp5 regulate circulating sugars and lipids

Levels of circulating nutrients are critical to starving cells. To explore if dIlp2, 3 and 5 regulate blood sugar levels in starving animals, we measured the trehalose content in the haemolymph. Like reported, feeding Δ*dIlp3* have higher sugar levels compared to controls confirming the sensitivity of our assay(Kim and Neufeld, 2015). In contrast, *dIlp2* and *dIlp5* mutants showed reduced circulating trehalose levels (Figure 2A). After 1 day starvation (short-term) trehalose levels in all genotypes dropped dramatically. Interestingly, after 5 days of nutrient depletion, sugar levels in Δ*dIlp2* recovered to levels of animals kept on food. In stark contrast, in Δ*dIlp5* trehalose levels dropped almost to levels beyond our detection limit displaying substantial differences in the sugar management between *dIlp2* and *dIlp5* mutants.

**Figure 2.**
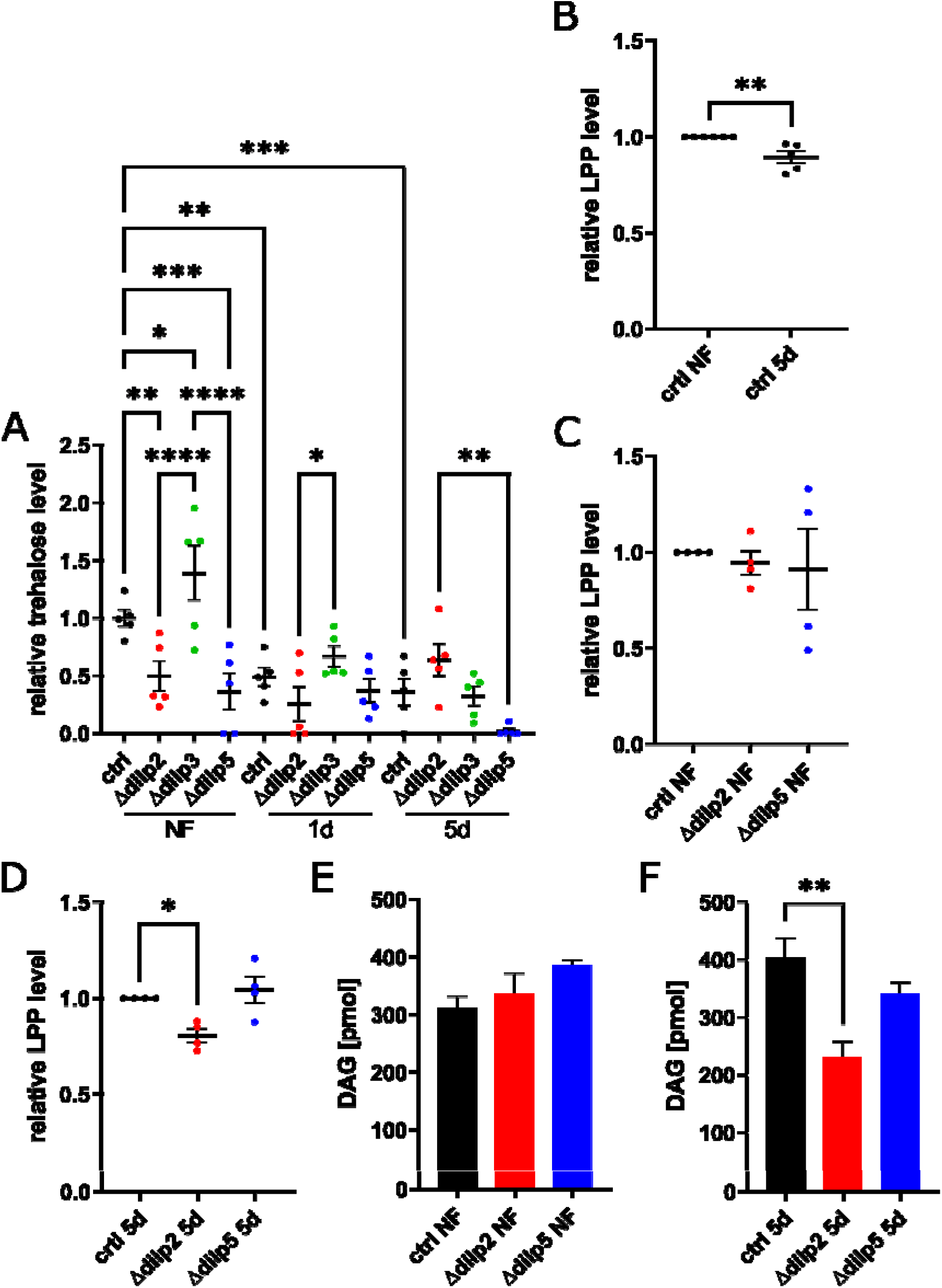
In starving animals lipid transport depends on dIlp2 and sugar homeostasis on dIlp5. (A) Shown is the relative trehalose level in the hemolymph of flies reared on normal food (NF) or starved for one (1d) or five (5d) days. Shown is the mean±SEM, one- way ANOVA followed by Fisher’s LSD multiple comparisons test (* p < 0.05, ** p < 0.01, *** p < 0.001, **** p < 0.0001). (B, C, D) Shown is the quantification of the relative LPP abundance in hemolymph detected by Western blotting of flies feeding on normal food (NF) or starved for five days (5d). Shown is the mean±SEM, unpaired Student’s t-test (** p < 0.01) (B) and one-way ANOVA followed by Tukey’s multiple comparisons test (* p < 0.05) (C, D). (E, F) Shown is the DAG content in fly heads measure by mass spectrometry of flies feeding on normal food (NF) (E) or starved for five days (5d) (F). Shown is the mean±SEM, one-way ANOVA followed by Tukey’s multiple comparisons test (* p < 0.05, ** p < 0.01).

We did not see differences in circulating protein levels in starving flies (Figure S1). However, to endure prolonged starvation fruit flies can mobilize fat resources and release lipids bound to lipophorin (LPP) into circulation (Palm et al., 2012). Some DILPs are suggested to control lipid homeostasis (Grönke et al., 2010; Semaniuk et al., 2018) and we decided to measure circulating lipoprotein particles from haemolymph by western blotting. We found that LPP levels drop in starving control flies (Figure 2B). We could not detect a difference in the amount of LPP in fed flies for mutant and control flies (Figure 2C). Starving *dIlp2* mutants had less LPP in circulation with respect to Δ*dIlp5* mutant or controls (Figure 2D). To measure the lipid load of LPP particles, we analysed diacylglyceride (DAG) levels in head samples from adult females by mass spectrometry. DAG is the preferred lipid load of lipoprotein particles in circulation in insects (Pennington and Wells, 2002), while cellular DAG levels represent only minority reports (Kennerly, 1987). Fed flies showed no differences in total DAG levels between tested genotypes (Figure 2E). In contrast, food deprived *dIlp2* mutants showed reduced total DAG levels if compared to Δ*dIlp5* or controls, possibly reflecting the lower amounts of LPP particles in hemolymph (Figure 2F). There appeared to be a general increase in fatty acid chain length and proportional decrease of saturated or mono-unsaturated DAG species in starving animals (Figure S2A-J). Both mutants, Δ*dIlp2* and Δ*dIlp5*, showed similar changes in their DAG profiles indicating that fatty acid qualities are not instructive for the lipid load onto LPP particles.

Taken together, the regulation of circulating sugars and lipids in starving flies is modulated by dIlp2 and dIlp5. In that context, both hormones seem to have different roles in maintaining nutritional homeostasis in food deprived *Drosophila*.

### dIlp2 and dIlp5 regulate the organization and activity of fat body organelles

In *Drosophila*, the fat body tissue represents vertebrate hepatocytes and adipocytes. Thus, the activity of fat body cells is likely essential in the nutritional regulation of starving flies. We have shown that *dIlp2* mutants produce less LPP particles, and have less lipids in circulation. We therefore speculated that the organization of lipid droplets (LDs) in the fat body will be different in starving Δ*dIlp2* with respect to controls. To visualize LDs, we stained fat body cells with BODIPY^505/515^ and imaged samples using fluorescent microscopy. Fed wild type flies had relatively small LDs dispersed throughout cells, whereas *dIlp2* and *dIlp5* mutants showed larger LDs (Figure 3A, B, C). In starving wild type animals, LDs increase with respect to fed controls (Figure 3A’). Further, LDs in starving Δ*dIlp2* appeared to be smaller than wild type controls, and in Δ*dIlp5* LDs were enlarged with respect to all other tested genotypes (Figure 3B’, C’). If the morphology of LDs in *dIlp5* mutants is determined by TAGs than we would expect to measure high lipid levels in these starving mutants. To validate that LD size and number correlate with lipid levels stored in these organelles, we decided to measure TAG levels by mass spectrometry from heads of adult mated females. We found no differences between fed Δ*dIlp2* and Δ*dIlp5* (Figure 3D). However, both mutants have less total TAGs stored with respect to controls. Thus, LD organisation cannot be directly translated into TAG levels. In starving wild type animals, TAG levels appear to increase with respect to fed siblings while in starving Δ*dIlp2* and Δ*dIlp5* TAG levels drop (Figure 3E). Hence, the lack of both, DILP2 or DILP5 promote the degradation of TAG in food deprived animals. Nevertheless, multivariant analyses of lipidomics data showed clear separation for the lipid species from starving Δ*dIlp2* and Δ*dIlp5* (Figure 3F). Thus, these mutants regulate many lipids in different ways and are therefore not redundant in their lipid turnover.

**Figure 3.**
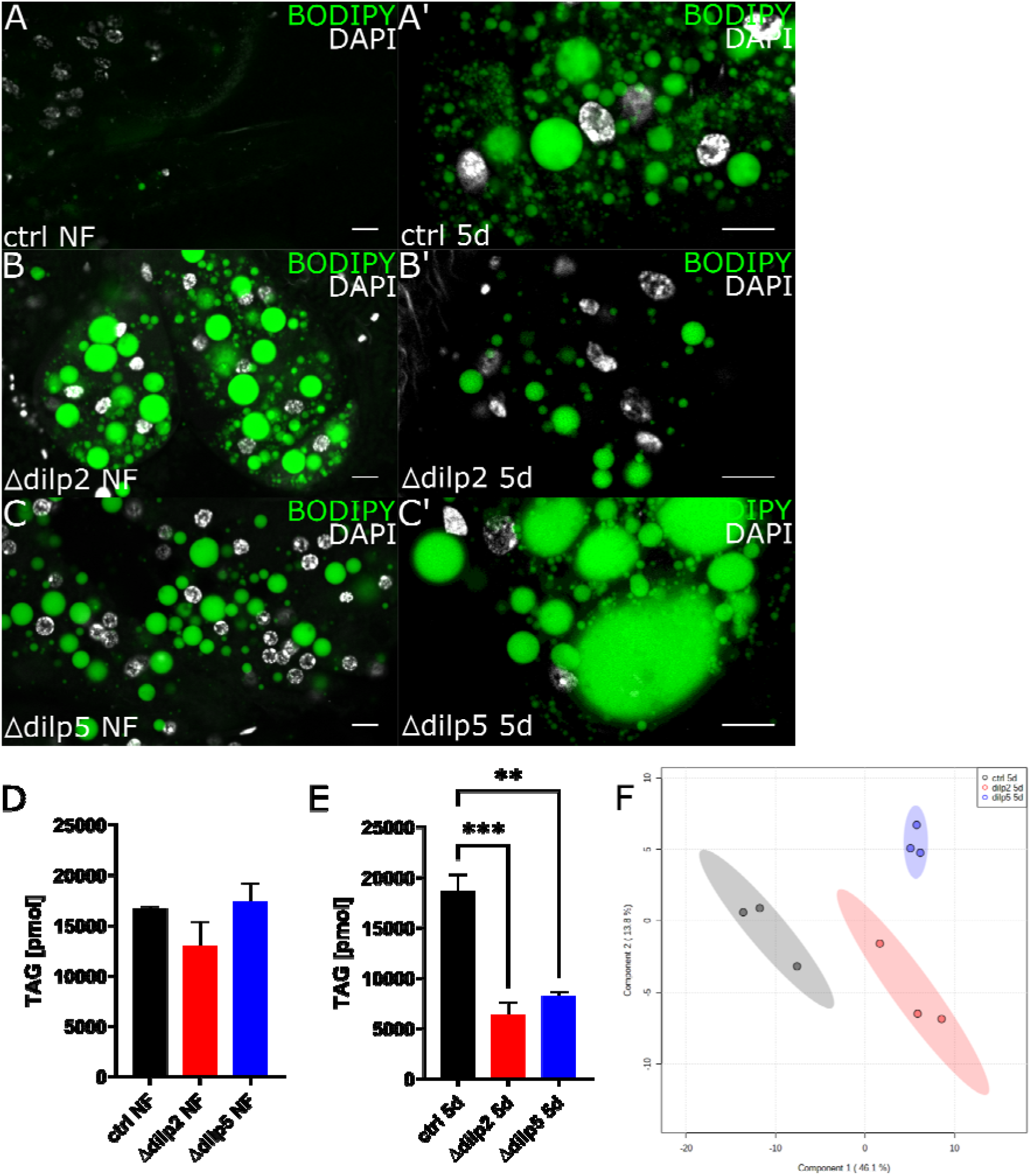
Organisation of lipid droplets and TAG storage is dependent in dIlp2 and dIlp5. (A-C) Microscopy pictures show exemplary images of fat body tissue of flies feeding on normal food (NF) (A, B, C) or starved for five days (5d) (A’-C’). Lipid droplets are stained with BODIPY505/515 and nuclei stained with DAPI. Scale bars indicate 10μm. (D, E) Shown is the TAG content in fly heads measure by mass spectrometry of flies feeding on normal food (NF) (D) or starved for five days (5d) (E). Shown is the mean±SEM, one-way ANOVA followed by Tukey’s multiple comparisons test (** p < 0.01, *** p < 0.001), n=3. (F) Shown is a score plot from PLS-DA analysis of mass spectrometry derived lipidomics data performed using MetaboAnalyst web-based platform.

Starvation shifted the ratios of TAG species (Figure S3A-J). We noticed that TAGs, composed of saturated and mono-unsaturated fatty acids, appeared to be more degraded compared to TAGs with poly-unsaturated fatty acids. In addition, we found that TAG species with longer fatty acid chain lengths accumulated in starving control and Δ*dIlp5* mutant flies. In contrast, starving *dIlp2* mutants showed no such proportional changes of TAG species.

These results correlate with earlier findings on DAG profiles in the haemolymph. The saturation profiles of circulating DAGs and LD stored TAGs could indicate that the recorded changes in TAG profiles are founded in the selective degradation of fatty acids.

### In starving animals dIlp2 or dIlp5 do not regulate Mitochondria in Fat body cells

In starving animals, hepatocytes are able to provide hepatic glucose by degrading glycogen to stabilize blood glucose levels. In *Drosophila* similar findings were made about S2 cells that degrade stored glycogen in response to DILP signalling (Post et al., 2018). We speculated, that in starving flies, fatty acids are degraded in mitochondria and resulting metabolites effecting hemolymph sugar levels. If true, we would expect differences in mitochondrial activity of starving Δ*dIlp2* with respect to Δ*dIlp5*.

To visualize mitochondria, we decided to stain live cells with MitoTracker – a dye that intercalates into mitochondrial membranes and reports to some extend mitochondrial activity. To test the assay, we targeted carnitine palmitoyltransferase 2 (*lpp>cpt2*^*RNAi*^) in fat body cells. The knock down of CPT2 prevents mitochondrial access to fatty acids reducing mitochondrial activity strongly. In parallel, we decided to knock down the microsomal triglyceride transfer protein (*lpp>mtp*^*RNAi*^). The loss of MTP in fat body cells prevents the loading of lipids onto lipoprotein particles and therefore, does not allow the clearance of fatty acids from fat body cells by secretion. Instead, we hoped that the surplus of fatty acids will be degraded by mitochondria intensifying the activity of these organelles. We found that MitoTracker hardly stains mitochondria in *lpp>cpt2*^*RNAi*^, while the reduction of MTP induced strong signals in respect to genetic controls (Figure S4). Utilizing our approach on fat body cells from fed flies showed a high mitochondrial activity in controls if compared to Δ*dIlp2* and Δ*dIlp5* (Figure 4A-C). Starving wild type flies showed reduced MitoTracker signals and were indifferent from the mitochondrial staining of both *dIlp* mutants (Figure 4A’-C’).

**Figure 4.**
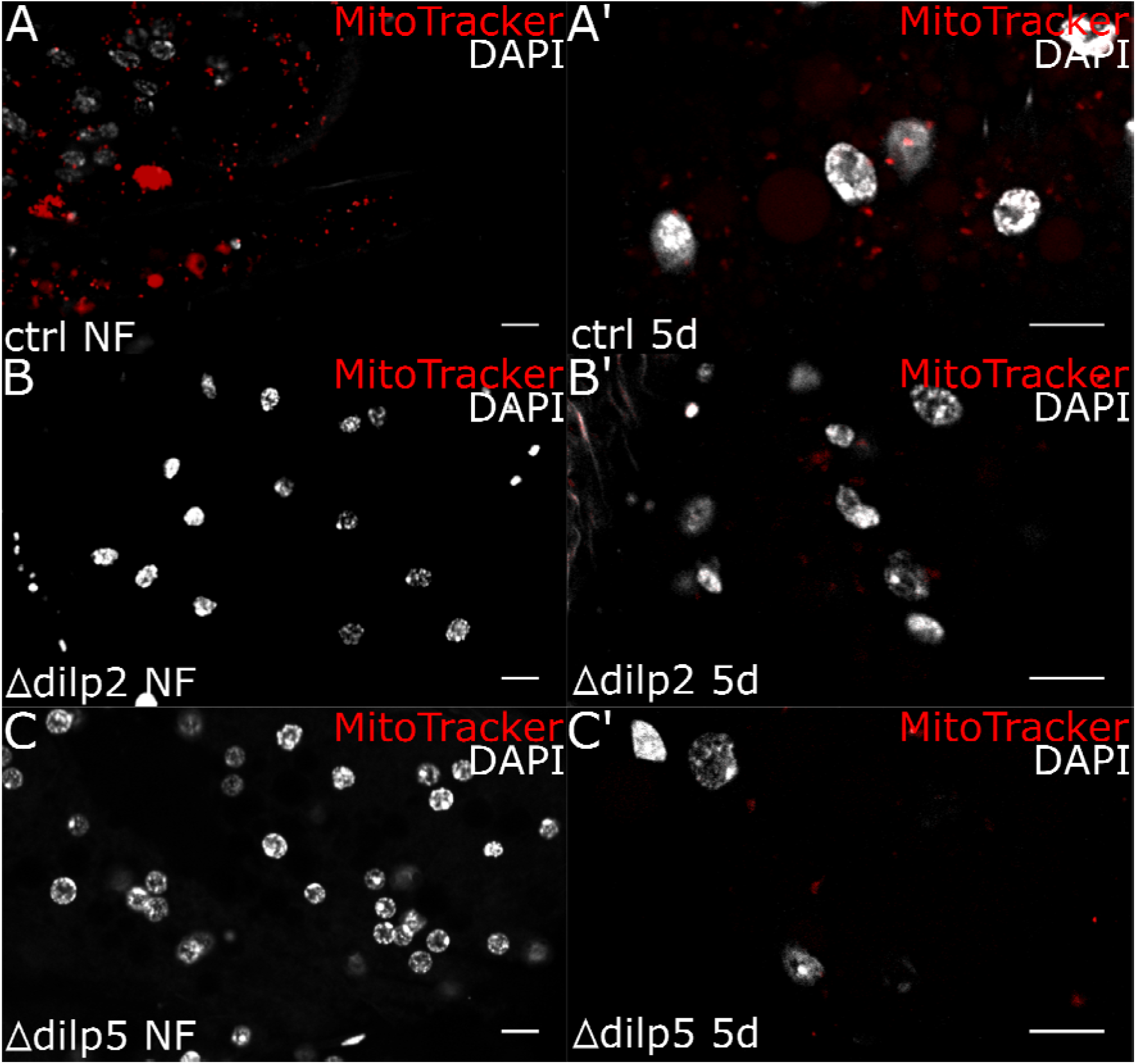
*dIlp2* and *dIlp5* mutants show low mitochondrial activity in fat body cells. (A- C) Microscopy pictures show exemplary images of fat body tissue of flies feeding on normal food (NF) (A-C) or starved for five days (5d) (A’-C’). Mitochondria are stained with MitoTracker and nuclei stained with DAPI. Scale bars indicate 10μm.

Nevertheless, the remaining mitochondrial gluconeogenesis in fat body cells could represent one way to buffer sugar levels in the haemolymph. To test this idea, we measured the haemolymph levels of starving controls, *lpp>cpt2*^*RNAi*^ and *lpp>mtp*^*RNAi*^ animals. We found no changes in the circulating sugar levels indicating that dIlp5 is not acting on fat body mitochondria, but is required elsewhere to buffer circulating trehalose levels in starving animals (Figure S4B). In addition, we wondered if dysfunctional mitochondrial beta oxidation of fatty acids or impaired lipid load of lipoprotein particles in fat body cells changes the amount of LPP present in blood system. We found no differences in LPP yields compared to controls (Figure S4C, D). Thus, the decrease of LPP particles in starving *dIlp2* mutants is possibly not based on the cellular utilization of fatty acids in fat body cells.

### dIlp5 signals over the Akt/PI3K pathway to regulate Akt^66^

Although DILP2 and DILP5 target the same InR, our data show that the lack of either produces different phenotypes in starving flies. For instance, fat body cells of starving Δ*dIlp5* show a different organization of LDs with respect to controls or Δ*dIlp2*. Lipid storage in LDs is dependent on insulin/insulin-like growth factor signaling (IIS), and we speculated that DILP2 possibly induces a different IIS activity than DILP5.

To investigate if DILP2 or DILP5 convey different IS signals, we assessed the phosphorylation state of Akt or MAPK by western blotting. We found that MAPK is less phosphorylated in starving control flies (Figure 5A). Further, we found no difference in the phosphorylation state of the MAPK in tested fed or starved animals from control flies (Figure 5B, C). Thus, the mitogenic branch of the IS cascade is unlikely affected by the absence of either DILP. Next, we examined the phosphorylation state at the amino acid Ser505 of both Akt isoforms, Akt^66^ and Akt^85^ using site-specific antibodies. We found that Akt^85^ is less phosphorylated at its Ser505 site in starving control flies (Figure 5D). In the mutants, Akt^85^ phosphorylation remained unchanged indicating that Akt^85^ is not affected by the absence of either DILP2 or DILP5 (Figure 5E, F). Further, we found that Akt^66^ is less phosphorylated at its Ser505 position in starving control flies (Figure 5G), however, we found that Akt^66^ phosphorylation is high in fed *dIlp5* mutants with respect to controls or Δ*dIlp2* (Figure 5H). However, in starving Δ*dIlp5* animals the phosphorylation of Akt^66^ decreased compared to controls (Figure 5I). Further, calculating total protein ratios between both Akt isoforms, we saw a change of Akt protein expression profiles for both *dIlp* mutants (Figure S5A, B). In addition, genetically induced expression of DILP5, but not DILP2 by IPC cells (*dIlp2>DILP2* and *dIlp2>DILP5*) elevated Akt^66^ phosphorylation levels under starvation only, indicating that DILP5 is required to potentiate Akt^66^ activity (Figure 5J and Figure S6A). Induced DILP5 expression did not affect the starvation resistance of adult *Drosophila*. In contrast, high DILP2 levels resulted in low survival rates under food deprivation indicating that high Akt^66^ activity is not affecting the starvation response (Figure S6B).

**Figure 5.**
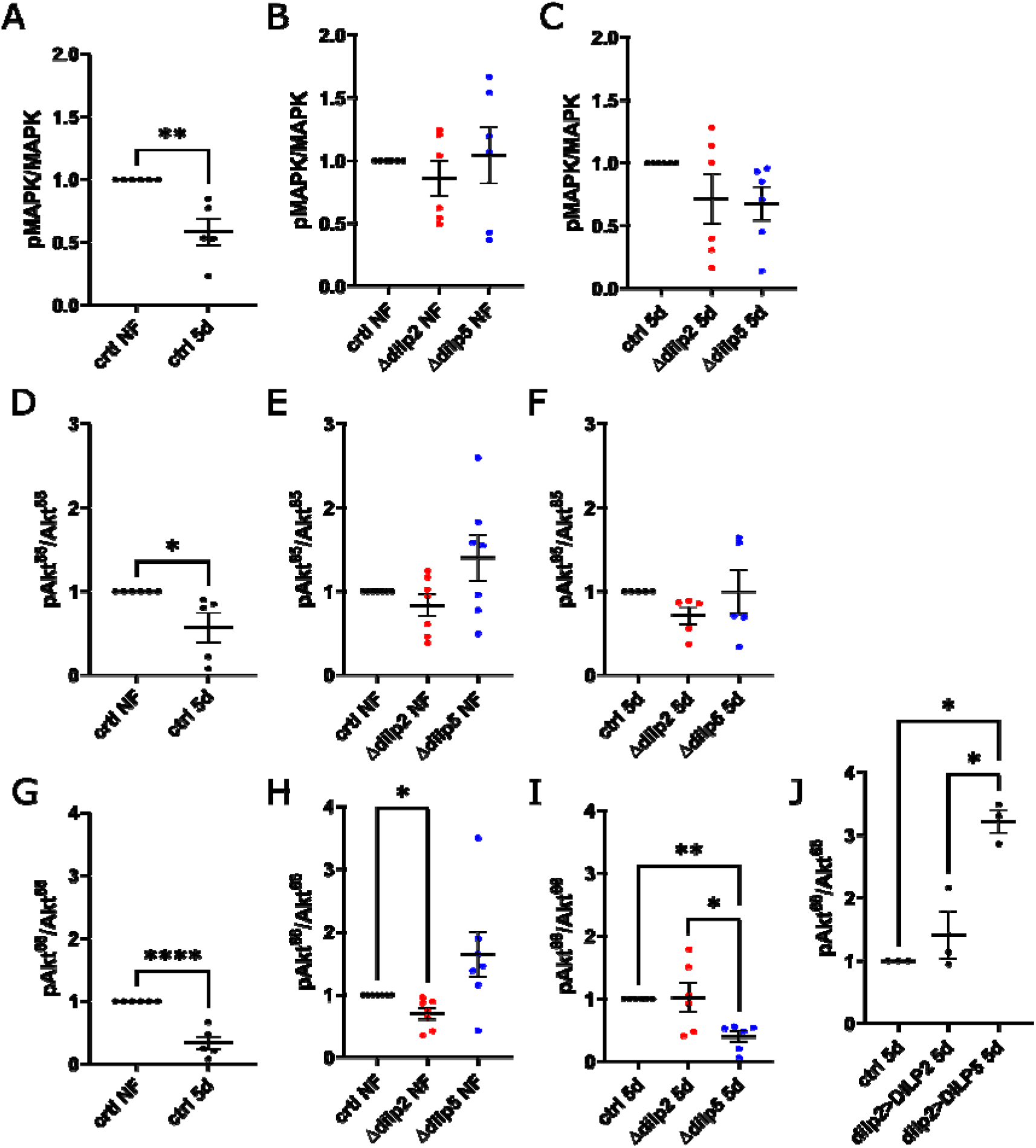
dIlp5 but not dIlp2 regulates Akt^66^ in starvation. (A-C) Shown is the quantification of phosphorylated MAPK relative to total MAPK detected by Western blotting. Signals are from fly heads of flies feeding on normal food (NF) or starved for five days (5d). Shown is the mean±SEM, unpaired Student’s t-test (** p < 0.01) (A) and one-way ANOVA followed by Tukey’s multiple comparisons test (B, C). (D-F) Shown is the quantification of phosphorylated Akt^85^ relative to total Akt^85^ detected by Western blotting. Signals are from fly heads of flies feeding on normal food (NF) or starved for five days (5d). Shown is the mean±SEM, unpaired Student’s t-test (* p < 0.05) (D) and one-way ANOVA followed by Tukey’s multiple comparisons test (E, F). (G-J) Shown is the quantification of phosphorylated Akt^66^ relative to total Akt^66^ detected by Western blotting. Signals are from fly heads of flies feeding on normal food (NF) or starved for five days (5d). Shown is the mean±SEM, unpaired Student’s t-test (**** p < 0.0001) (G) and one-way ANOVA followed by Tukey’s multiple comparisons test (* p < 0.05, ** p < 0.01) (H-J).

### PI3K activity in fat body cells modulates the starvation response

We found a regulation of Akt^66^ by dIlp5 but not dIlp2. To investigate, if the systemic dIlp signaling is mimicked by local IIS in the fat body, we decided to manipulate the PI3K activity in fat body cells. The ratio between PIP2 and PIP3 in cellular membranes is instructive for the recruitment of Akt (Manna and Jain, 2013). We decided to reduce PIP3 genetically in fat body cells by an RNAi-mediated knockdown of the enzyme PI3K (PI3K^KD^) and thereby affect the phosphorylation of Akt. To validate that the knock down of PI3K is not altering the mitogenic branch of the insulin signal cascade, we decided to assess the phosphorylation state of the MAPK. As expected, MAPK activity was independent from PI3K manipulations (Figure S7A). Next, we probed for the phosphorylation of Akt isoforms in PI3K^KD^ flies. We found that depletion of PI3K in starving animals did not affect the phosphorylation profile at the Akt^85Ser505^ position (Figure 6A). In stark contrast, the PI3K^KD^ reduced Akt^66^ phosphorylation at its Akt^66Ser505^ position in starving animals (Figure 6B). However, PI3K^KD^ do not change the ratio between Akt^85^ and Akt^66^ (Figure S7B).

**Figure 6.**
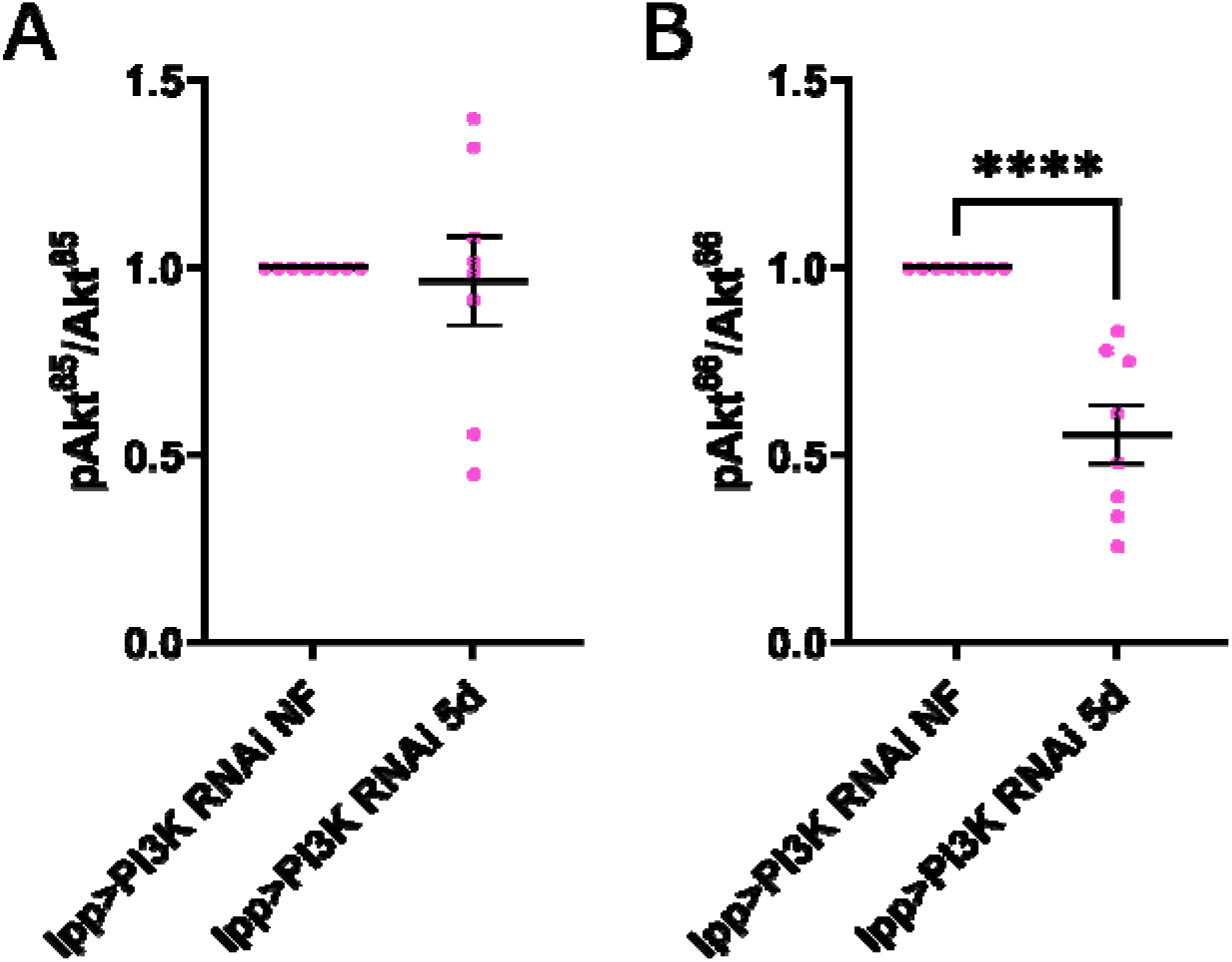
PI3K activity in fat body cells regulates Akt^66^ phosphorylation. (A, B) Shown is the quantification of phosphorylated Akt^85^ relative to total Akt^85^ (A) or phosphorylated Akt^66^ relative to total Akt^66^ (B) detected by Western blotting. Signals are from fly heads of flies feeding on normal food (NF) or starved for five days (5d). Shown is the mean±SEM, unpaired Student’s t-test (**** p < 0.0001).

With the PI3K knockdown in fat body cells, we hoped to modify IIS and assess the role of dIlps in lipid mobilization. The dampened IIS in PI3K^KD^ flies could lead to impaired build-up of fat stores. We found that measured TAG levels of fed PIK3^KD^ were not different from controls (Figure 7A) but starving PIK3^KD^ have low TAG yields with respect to controls (Figure 7B). Despite TAG levels are lower during starvation, similar to *dIlp* mutants, PI3K^KD^ displayed similar starvation resistance to control flies (Figure S7C). At cellular level, in fat body cells mitochondrial activity was reduced in fed or starving PI3K^KD^ flies (Figure 6C, C’) with respect to controls. We found similar enrichments of LDs in fed and starved PI3K^KD^ animals (Figure 6D, D’). Next, we tested for LPP levels present in the haemolymph. We found no differences between fed and starving PI3K^KD^ animals, and LPP yields were similar to respective controls (Figure 6E, F).

**Figure 7.**
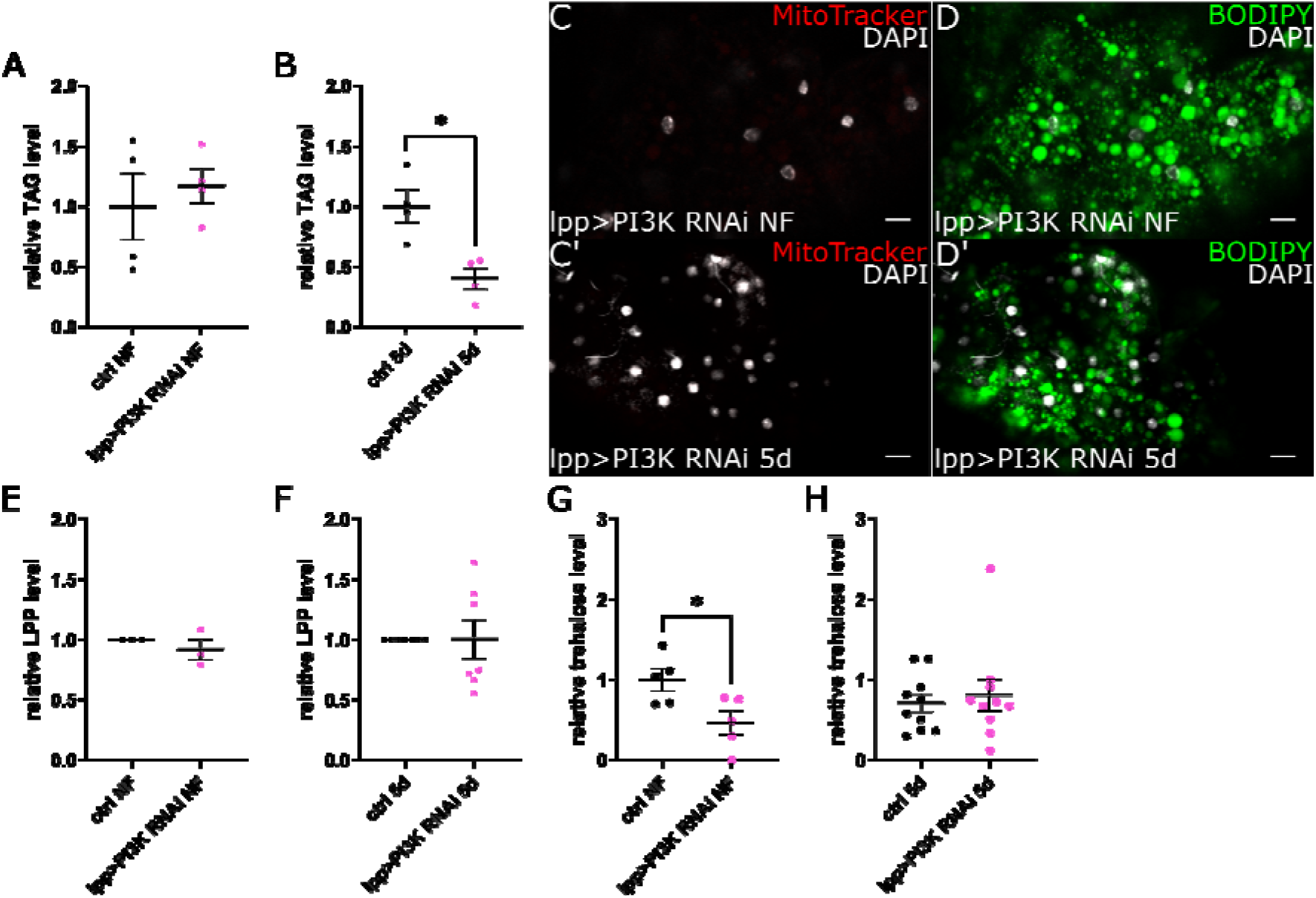
PI3K activity in fat body cells modulates lipid droplet morphology and TAG turnover. (A, B) Shown is the quantification of the relative TAG abundance detected by thin- layer chromotography in fly heads from flies feeding on normal food (NF) (A) or starved for five days (5d) (B). Shown is the mean±SEM, unpaired Student’s t-test (* p < 0.05). (C, D) Microscopy pictures show exemplary images of fat body tissue of flies feeding on normal food (NF) (C, D) or starved for five days (5d) (C’, D’). Lipid droplets are stained with BODIPY505/515, mitochondria are stained with MitoTracker and nuclei stained with DAPI. Scale bars indicate 10μm. (E, F) Shown is the quantification of the relative LPP abundance in hemolymph detected by Western blotting of flies feeding on normal food (NF) (E) or starved for five days (5d) (F). Shown is the mean±SEM, unpaired Student’s t-test. (G, H) Shown is the relative trehalose level in the hemolymph of flies reared on normal food (NF) or starved for five days (5d). Shown is the mean±SEM, unpaired Student’s t-test (* p < 0.05).

We found that in prolonged starving animals PI3K regulates the activity of Akt66 and hence, likely the activity of the transcription factor FOXO. The activity of FOXO is directly linked with the lipid but also blood-sugar levels(Lee and Dong, 2017). We found that PI3K^KD^ kept on food had low circulating sugar levels with respect to controls (Figure 6G) and sugar levels in starving PI3K^KD^ were similar to controls (Figure 6H).

Taken together, we have shown that PI3K^KD^ flies mimic cellular metabolic responses of each of the *dIlp* mutant to prolonged starvation but differ in their starvation resistance and LD organisation of fat body cells. Interestingly, PI3K activity is not required to build TAG stores and starving *dIlp* mutants and PIK3^KD^ consume stored TAGs faster than controls.

## Discussion

In starving *Drosophila*, the non-canonical dIlp6 peptide is essential to increase starvation resistance (Chatterjee et al., 2014). More recently, significant roles for DILP2 and DILP5 were proposed by regulating the glycogen phosphorylase required to mobilize energy from glycogen stores (Post et al., 2018). Here we show that dIlp2 and dIlp5 are instrumental for the lipid metabolism of starving adult *Drosophila*. We found that dIlp2 regulates the release of lipoprotein particles required to transport lipids in the blood system, and dIlp5 modulates the morphology and load of cellular lipid droplets (LDs).

In starving mammals, cellular membrane lipids are degraded, and in the process released FAs are bound as TAGs. We found evidence that food deprived *Drosophila* use the same principles to build initial TAG stores. We show that TAG levels in starving fruit flies increase and the poly-unsaturated fatty acids (PUFA) proportion raises in the hemolymph. Flies encode three different L9 desaturases and hence, are limited in their FA production to saturated fatty acids (SFAs) and mono/unsaturated fatty acids (MUFAs) (Fang et al., 2009). Thus, the increase of PUFAs in the hemolymph excludes FA *de novo* synthesis and point to phospholipids as the most likely source of FAs. On the other hand, starving animals show a tendency to degrade preferentially SFAs to produce energy. The selective retention of long MUFAs appears to be a highly conserved mechanism and is shown for food deprived birds or hibernating mammals (Geiser, 2021; Twining et al., 2016).

Animals use different lipoprotein particles to transport lipids in the circulatory system(Palm et al., 2012). In *Drosophila*, the fat body derived LPP is the main lipid carrier (Canavoso et al., 2001) and we found a reduction of LPP particles in starving flies. Interestingly, the amount of lipids loaded onto LPP was higher than in fed controls suggesting that starving animals produce large LPP particles. The expression of lipoproteins is regulated by FOXO. FOXO represents the terminal end of the cellular insulin signal cascade and is active in hungry animals. In addition, the formation of TAG stores in starving animals is part of the anabolic metabolism. Thus, we speculated that insulin peptides regulate lipid turnover in food deprived flies.

Adult flies produce multiple neuronal DILP peptides, but downregulate the expression of many entering malnutrition. The notable exception is DILP2, but its functional role is somewhat less clear (Bai et al., 2012; Grönke et al., 2010; Ikeya et al., 2002; Sudhakar et al., 2020). We found that the loss or gain of DILP2 increases the starvation sensitivity of adult *Drosophila* pointing to the necessity to keep the functional DILP2 expression upon nutritional deprivation. Investigating nutritional cues in circulation of fed animals, we confirmed that DILP3 clears sugars in the circulatory system (Kim and Neufeld, 2015). In contrast, *dIlp2* and *dIlp5* mutants displayed lower blood-sugar levels with respect to genetic controls. Our results for starvation resistance sidelined the possible involvement of dIlp3 in our study, especially since *dIlp2, 3* double mutants did not increase the phenotype of Δ*dIlp2* or Δ*dIlp5* flies. We found that starving flies lacking dIlp2 but not dIlp5 have less LPP and transport less lipids. Thus, continuous dIlp2 expression is required to stimulate LPP expression in fat body cells. Δ*dIlp5* show circulating LPP and lipid yields comparable to controls. However, the loss of Δ*dIlp5* induces an increase in the size of LDs of fat body cells.

In contrast, *dIlp2* mutants show small and less LDs. This result is unexpected since TAG levels in starving Δ*dIlp2* and Δ*dIlp5* are low with respect to controls.

The nature of the cargo filling the LDs of Δ*dIlp5* remains to be identified; however, we speculate that a disproportional increase of sterol esters could offer a possible solution. If true, starving Δ*dIlp5* flies maintain a different sterol homeostasis than Δ*dIlp2* or controls. In addition, dIlp5 potentially initiates different insulin signaling in fat body cells than dIlp2. To test the latter idea, we probed the phosphorylation status of the enzyme Akt, positioned downstream of the InR in the cellular signal cascade. Different Akt isoforms are expressed in *Drosophila* and in contrast to the well characterized Akt proteins in mammals (Andjelkovic et al., 1995; Gonzalez and McGraw, 2009), it remains unclear if each individual protein isoform fills a specific role in insulin signaling. We found that in starving adult animals both expressed Akt isoforms, Akt^66^ and ^85^, are less phosphorylated at their Ser505 position with respect to fed controls. We found that neither dIlp2 nor dIlp5 regulate Akt^85^ in fed animals. Instead, we show that in animals kept on food, the phosphorylation of Akt^66^ is dependent on dIlp2 signaling; however, in starving animals dIlp5 and not dIlp2 signaling regulates Akt^66^ phosphorylation. This switch in metabolic regulation between dIlp2 and dIlp5 after nutritional deprivation remained elusive so far. We speculated that specific cell types respond to nutritional hardships with radical functional changes. Thus, we focused on fat body cells that are known to regulate starvation stress responses. To manipulate Akt signaling we decided to study fat body specific knock downs of PI3K. Like expected, the loss of PI3K results in low Akt^66^ phosphorylation levels in starving animals. Indeed, low Akt^66^ activity correlated in both genetic background in the induction of LDs. It remains unclear if LDs are directly regulated by Akt^66^ or if the activity of the canonical Akt target FOXO modulates the dynamic behaviors of LDs.

The question that remains is how specificity of dIlp2 and dIlp5 signaling can be produced through one common receptor. It was shown for *Drosophila* S2 cells that DILP2 and DILP5 produce similar transcriptional patterns while inducing different phosphorylation kinetics in the IIS (Post et al., 2018). Solutions offered to the problem include variations in InR affinity, spatiotemporal concentration patterns of dIlps, yet undiscovered co-receptors and binding proteins, and differing phosphorylation patterns in the IIS cascade. Our study supports the idea that individual dIlps act downstream the insulin receptor by selectively activating different Akt isoforms leading to metabolic responses important to adapt to nutritional deprivation.

## Material and Methods

### Fly husbandry and lines

Fly stocks were raised at 20°C on normal food (NF) (Prince et al., 2021) in a day/night cycle. The following fly lines were used: *foxo[mCherry]* (#80565 from Bloomington *Drosophila* Stock Center (BDRC)), Δ*dIlp2* and Δ*dIlp5* (from S. Grönke), *dIlp2-Gal4* (#37516 from BDRC), *UAS-dIlp2* and *UAS-dIlp5* (from Eaton lab), *lpp-Gal4* (from Eaton lab), *UAS-cpt2 RNAi* (from Schirmeier lab), UAS-mtp RNAi (from Eaton lab), UAS-PI3K RNAi (from Knust lab).

Experimental flies were assayed after being reared on NF for 5 to 15 days. All experiments were conducted at 20°C. *foxo[mCherry]* flies were used as a genetic control (ctrl) for *dIlp* mutant experiments while *lpp-Gal4* x *foxo[mCherry]* was used as a control for experiments involving genotypes driven by lpp-Gal4 and *dIlp2-Gal4* x *foxo[mCherry]* for experiments with genotypes driven by *dIlp2- Gal4*.

### Survival assay

Flies (3 males and 9 females) were transferred to agar plates (1% agar, 0.4% nipagin) and alive flies were counted every 1-2 days. Data is represented as the percentage of alive flies per day. One Plates equals one replicate.

### Feeding behavior assay

Flies (3 males and 9 females) were transferred overnight on plates (1% agar, 0.4% nipagin). Agar plates were equipped with one NF source positioned in the middle of each plate. The next day, feeding behavior was recorded for 1h (1 frame per second). Food attraction was measured by flies attending the food measured by calculating the mean grey values in the area covering the food source. Values were normalized to the plain food background.

### Quantitative RT-PCR

Total RNA was isolated from 10 Female fly heads (*foxo[mCherry]*) according to the manufacturer protocol (RNeasy Mini Kit from QIAGEN). mRNA abundance of the *dIlp2* and *dIlp5* gene was calculated by the method of comparative C_T_ and normalized to each other. Used primers are published(Grönke et al., 2010).

### Hemolymph trehalose and protein quantification

To collect hemolymph, frozen female flies were thawed on ice for 1h. Thereafter, ice-cold PBS was added to samples. Samples were incubated for 5 min on ice and centrifuged with a benchtop centrifuge twice, then supernatant was collected in a fresh tube. Trehalose and protein were quantified according to manufacturer from Trehalose Kit (Megazyme) and BCA Kit (Pierce). Data was normalized to the control flies’ measurements feeding on normal food.

### Western blotting

Femal flies were snap-frozen in liquid nitrogen, three fly heads were pooled per sample. Samples were homogenized on ice in PBS, heat treated at 95°C for 10min and subsequently loaded onto Tris-SDS-PAGE. Proteins were transferred on membranes via tank blotting. Membranes were blocked in 5% BSA in 0.1% Triton X-100/PBS and subsequently probed with antibodies including anti-Akt (Invitrogen), anti-Akt^pSer505^ (Cell Signaling), anti-Akt^pThr308^ (Invitrogen), anti-MAPK (Cell Signaling), anti-pMAPK (Cell Signaling), anti-LPP (Eaton lab) and HRP conjugated secondary antibodies (Thermo Fisher). Detected signals were normalized to control flies’ measurements. Data was corrected for outliers using the IQR method.

### Mass spectrometry

Samples of female fly heads were collected according to protocols published by Lipotype (https://www.lipotype.com). Lipid extraction and measurements were performed by services offered by Lipotype.

### Thin-Layer Chromatography (TLC)

Heads of female flies were homogenized and lipids were extracted with 10:1 Methanol:Chloroform. Lipids were air-dried, stored at -80 °C for no longer than one night and subsequently resolved in 2:1 Methanol:Chloroform. Lipid extracts were loaded onto a silica plate (VWR) and separated with: Heptane:Diethylether:Acetic acid (70:30:1, v/v/v) or Chloroform:Methanol:water (75:25:2.5, v/v/v). Lipids were detected using primulin dye and plates were scanned with ImageQuant software. Neutral lipid signals were normalized to Phosphatidylethanolamine (PE) signal. Then, signals were normalized to control flies’ measurements.

### Microscopy

Female flies were dissected in ice-cold Grace’s Insect Medium, stained on ice with BODIPY505/515 (final conc. 0.5 μg/ml), MitoTracker (final conc. 100nM) and DAPI (final conc. 1 μg/ml) and subsequently imaged with confocal microscope (ZEISS-LSM 700).

### Data analyses and statistics

TLC, Western blots and microscopy images were anlaysed using FIJI ImageJ software. Lipidomics data were analyzed with MetaboAnalyst 5.0 (Pang et al., 2021). Graph depiction and statistical analyses was done using GraphPad Prism 9.

## Acknowledgements

We thank the light microscopy facility and Katharina Ganß for technical assistance, MG for critical comments.

LT/MB supported by FOR2682/TP3

