## Supplement Data for "Insulin regulates systemic lipid traffic in starving animals"

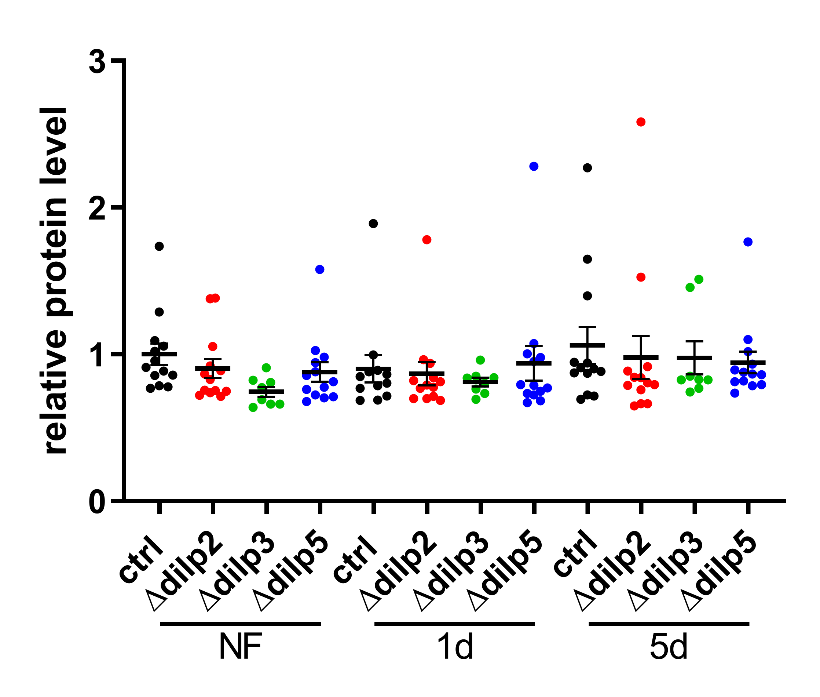

**Figure S1. Proteins in circulation are not regulated by neural dIlps.** Shown is the relative protein level in the hemolymph of flies reared on normal food (NF) or starved for one (1d) or five (5d) days. Shown is the mean±SEM, one-way ANOVA followed by Fisher’s LSD multiple comparisons test.

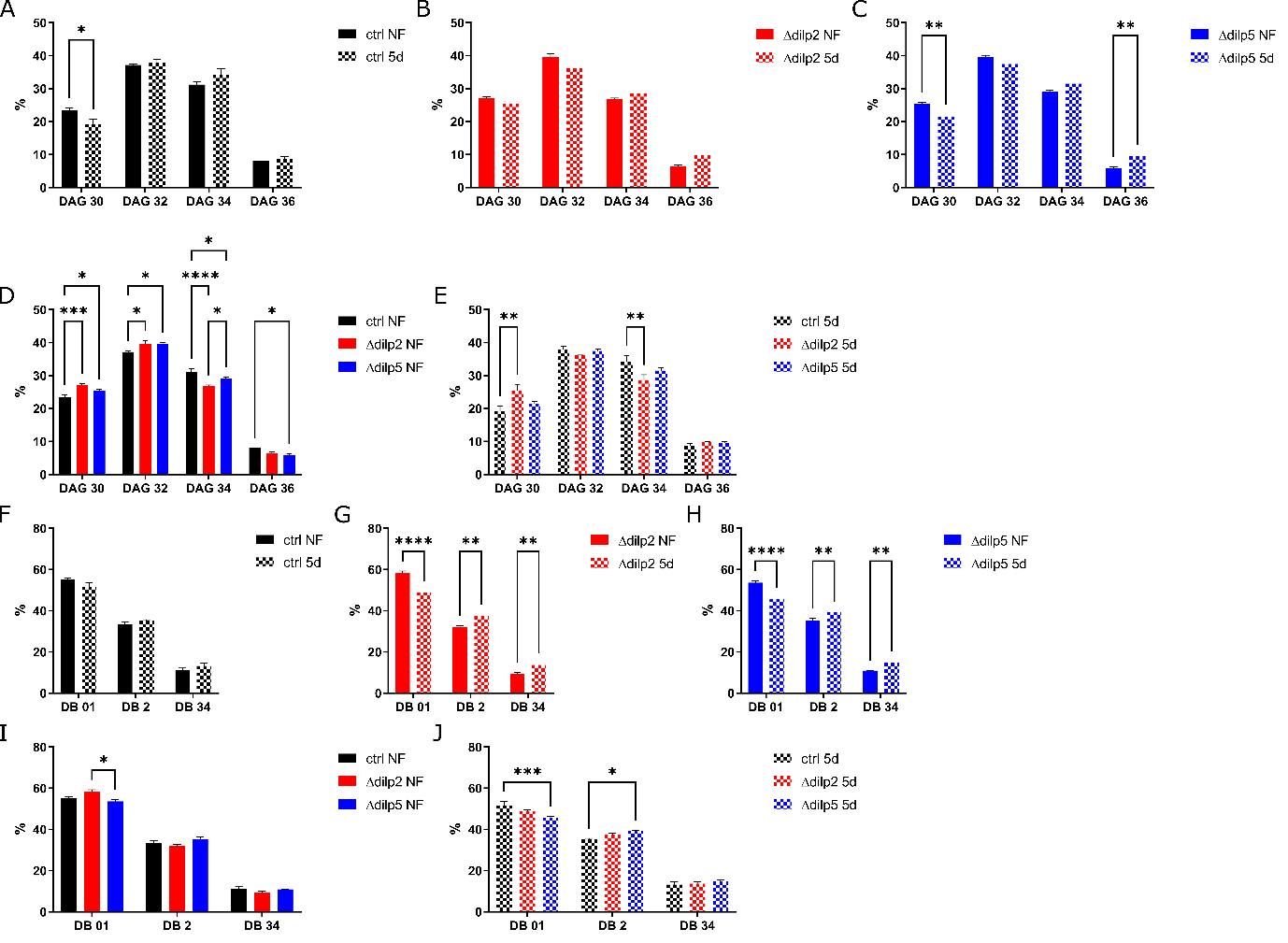

**Figure S2. DAG profiles in fed and starving animals.** (A-E) Shown is the distribution of DAG species, quantified as a sum of all species with the same carbon number, in total DAGs in percentage in animals feeding on normal food (NF) or starved for five days (5d). (F-J) Shown is the distribution of the number (0-4) of double bounds (DB) in total DAGs in percentage in animals feeding on normal food (NF) or starved for five days (5d). Shown is the mean±SEM, two-way ANOVA followed by Sidak’s multiple comparisons test (* p < 0.05, ** p < 0.01, *** p < 0.001, **** p < 0.0001), n=3.

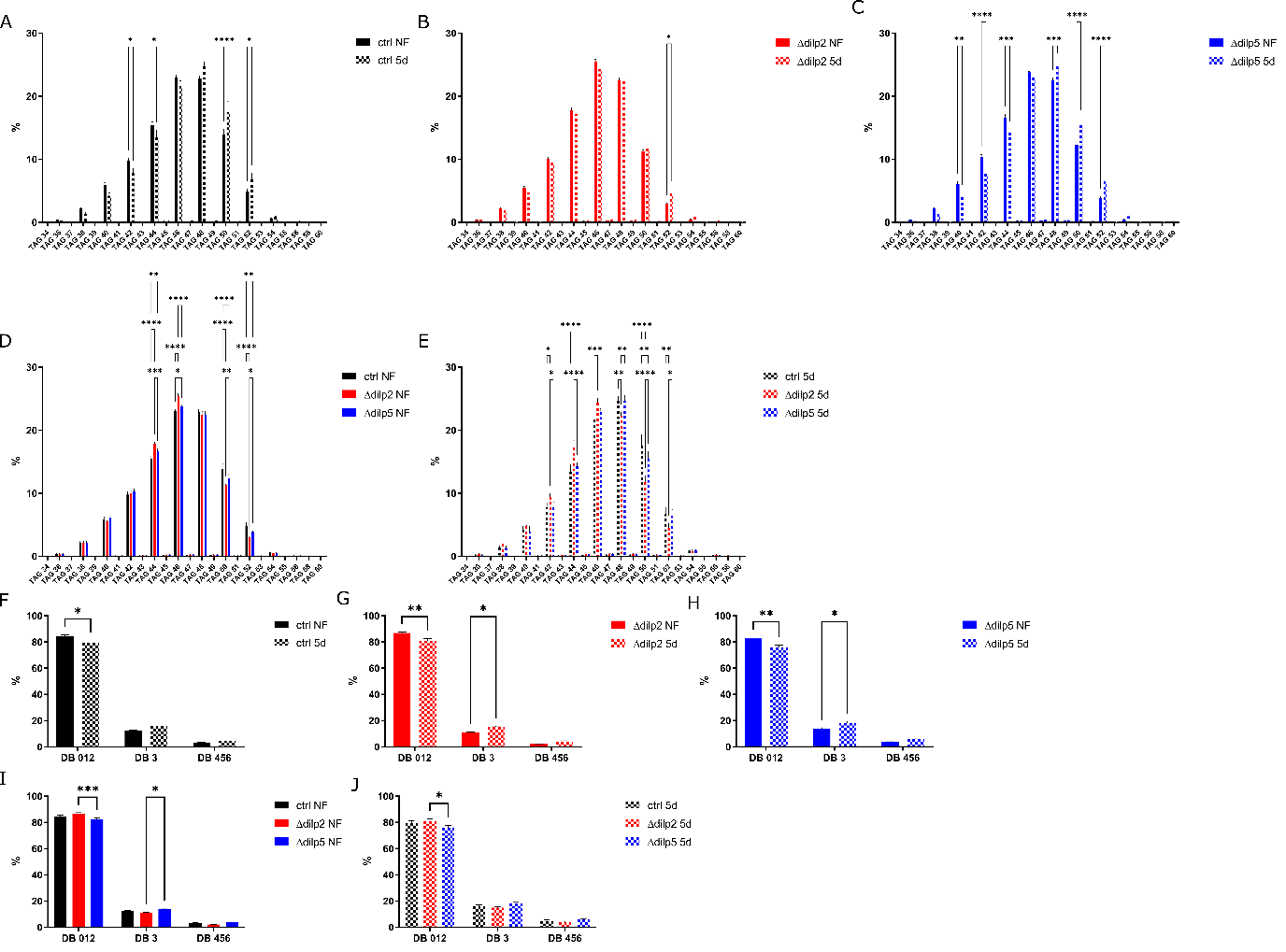

**Figure S3. TAG profiles in fed and starving animals.** (A-E) Shown is the distribution of TAG species, quantified as a sum of all species with the same carbon number, in total TAGs in percentage in animals feeding on normal food (NF) or starved for five days (5d). (F-J) Shown is the distribution of the number (0-6) of double bounds (DB) in total TAGs in percentage in animals feeding on normal food (NF) or starved for five days (5d). Shown is the mean±SEM, two-way ANOVA followed by Sidak’s multiple comparisons test (* p < 0.05, ** p < 0.01, *** p < 0.001, **** p < 0.0001), n=3.

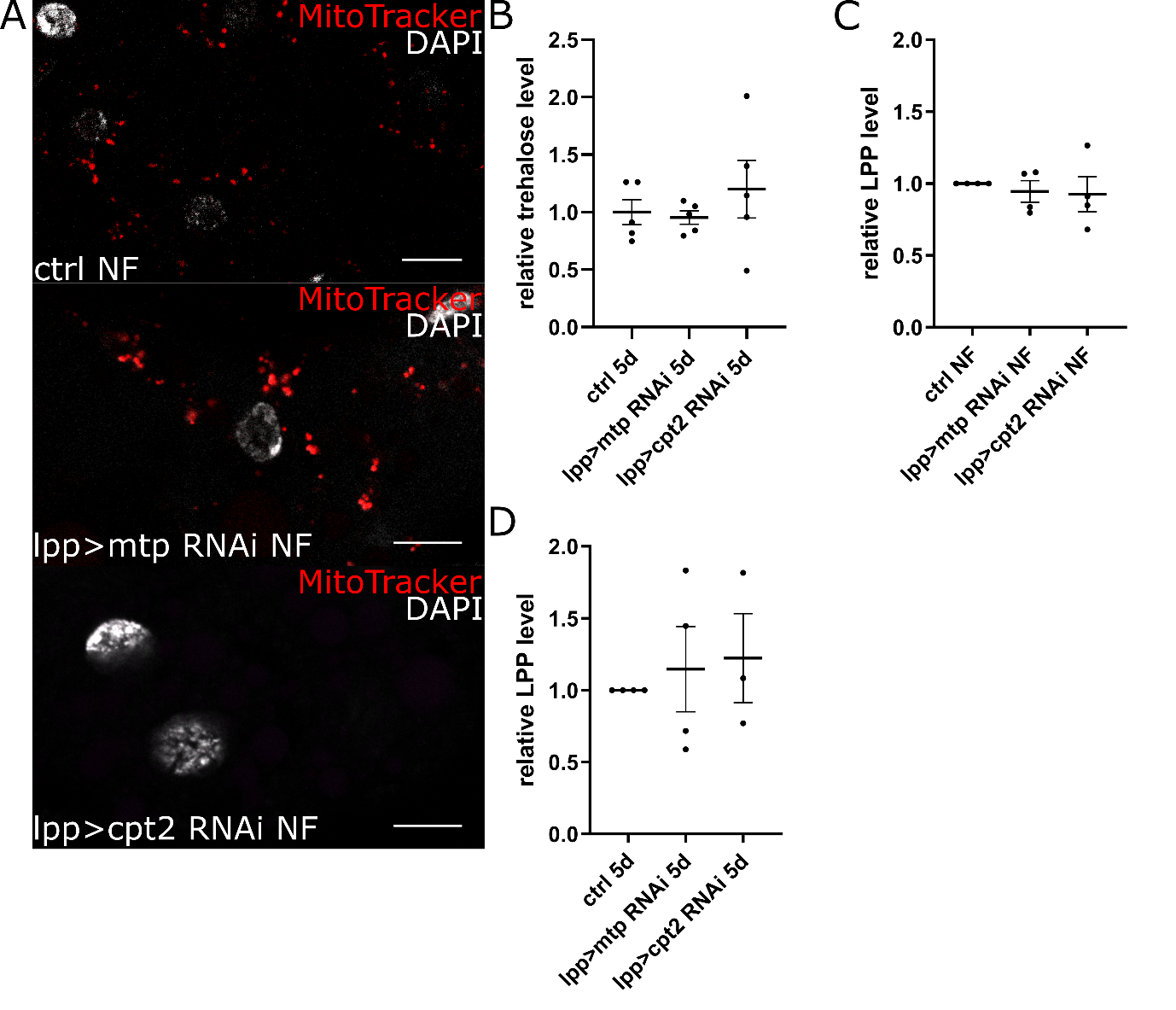

**Figure S4. Mitochondrial activity is dependent on cpt2 and mtp in fat body cells.** (A) Microscopy pictures show exemplary images of fat body tissue of flies feeding on normal food (NF). Mitochondria are stained with MitoTracker and nuclei stained with DAPI. Scale bars indicate 10µm. (B) Shown is the relative trehalose level in the hemolymph of flies starved for five days (5d). Shown is the mean±SEM, one-way ANOVA followed by Tukey’s multiple comparisons test. (C, D) Shown is the quantification of the relative LPP abundance in hemolymph detected by Western blotting of flies feeding on normal food (NF) (C) or starved for five days (5d) (D). Shown is the mean±SEM, one-way ANOVA followed by Tukey’s multiple comparisons test.

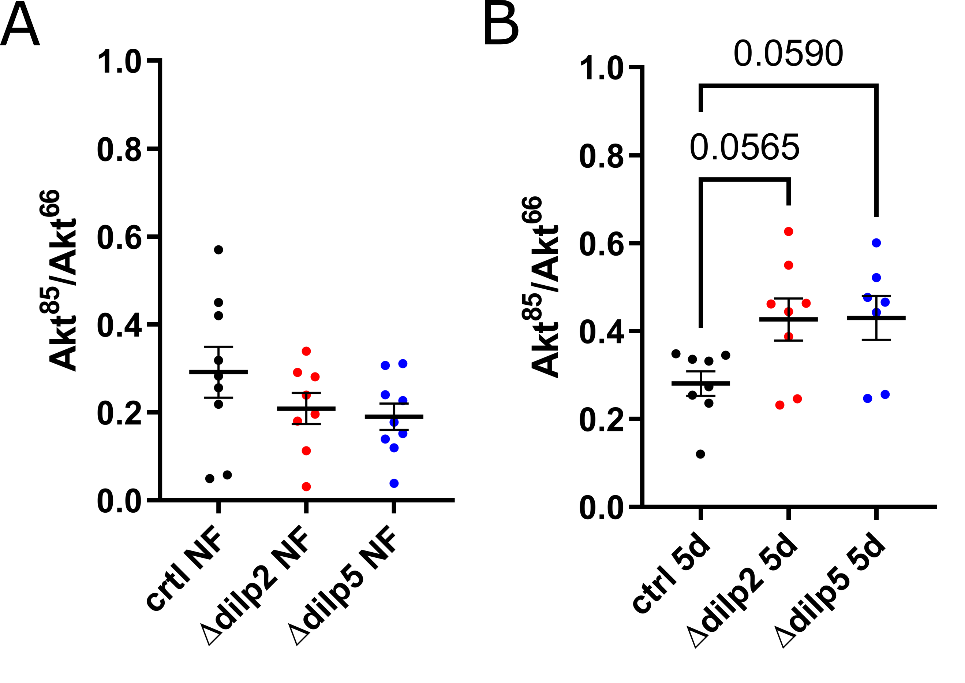

**Figure S5. Akt^85^ to Akt^66^ ratios in fed and starving animals.** (A, B) Shown is the quantification of Akt^85^ relative to Akt^66^ detected by Western blotting. Signals are from fly heads of flies feeding on normal food (NF) (A) or starved for five days (5d) (B). Shown is the mean±SEM, one-way ANOVA followed by Tukey’s multiple comparisons test.

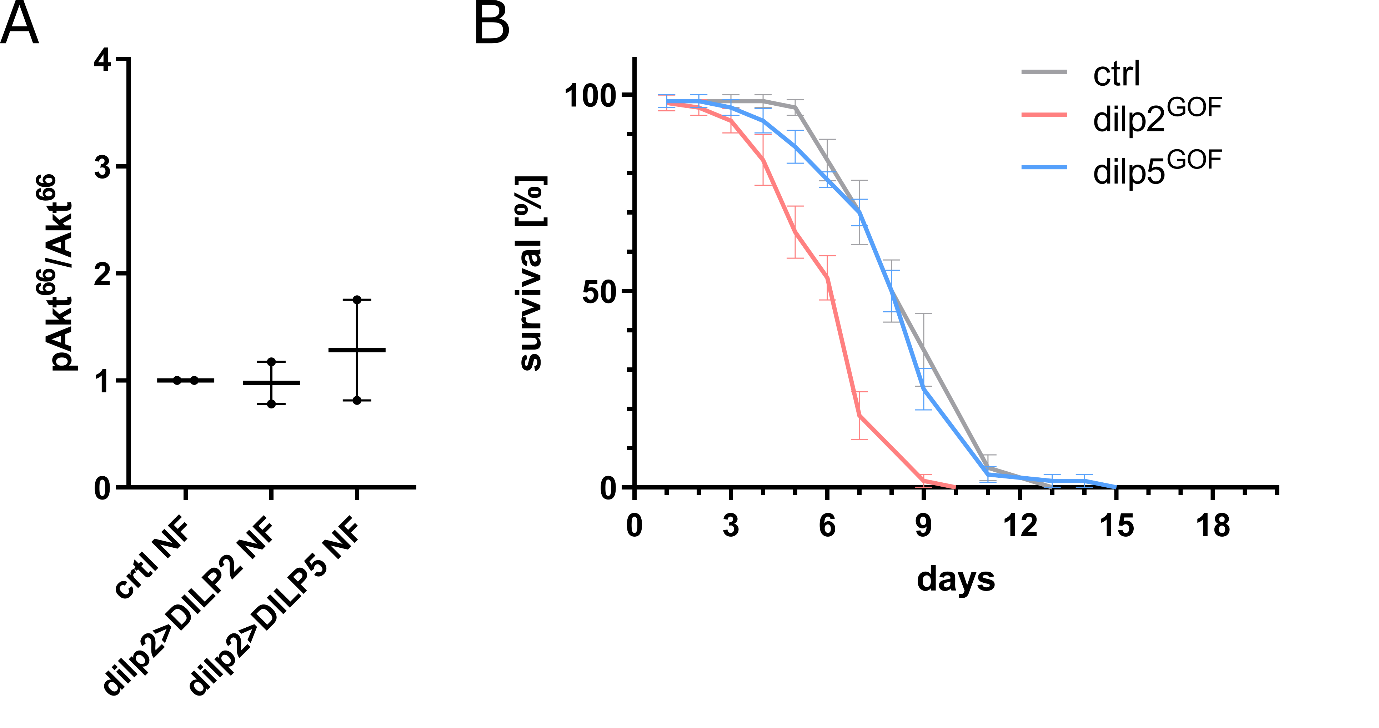

**Figure S6. *dIlp2* overexpression decreases starvation resistance.** (A) Shown is the quantification of phosphorylated Akt^66^ relative to total Akt^66^ detected by Western blotting. Signals are from fly heads of flies feeding on normal food (NF). Shown is the mean±SEM, ne-way ANOVA followed by Tukey’s multiple comparisons test. (B) Plotted is the survival curve of flies exposed to long-term starvation on agar plates (n = 5).

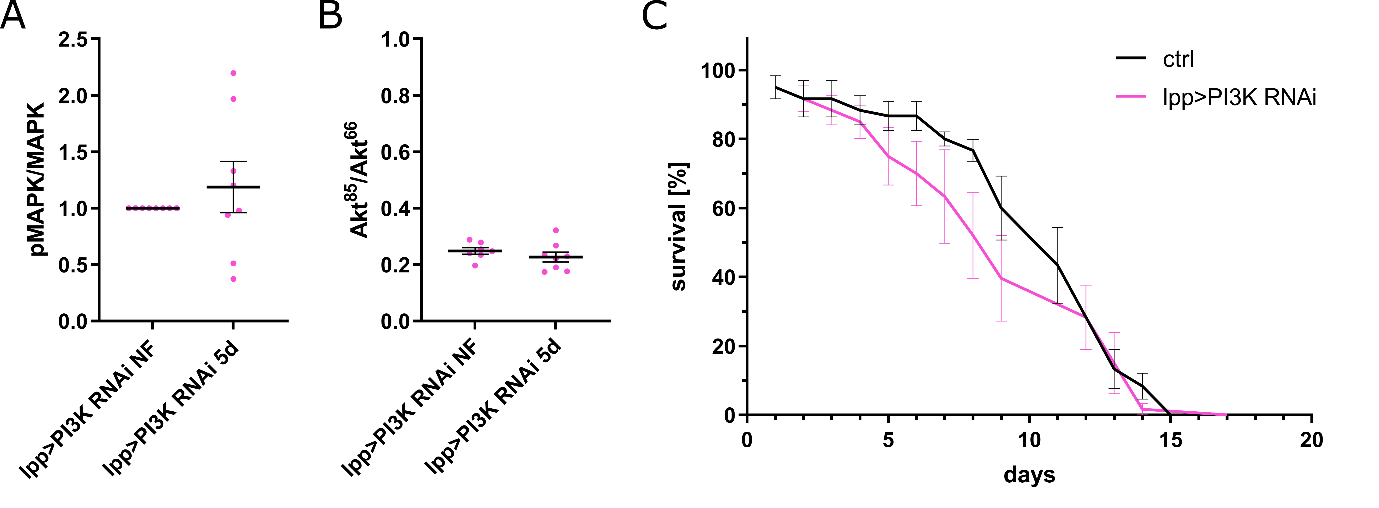

**Figure S7. Loss of PI3K activity in fat body cells is not decreasing starvation resistance.** (A, B) Shown is the quantification of phosphorylated MAPK to total MAPK (A) or phosphorylated Akt^85^ relative to total Akt^85^ (B) detected by Western blotting. Signals are from fly heads of flies feeding on normal food (NF) or starved for five days (5d). Shown is the mean±SEM, unpaired Student’s t-test. (C) Plotted is the survival curve of flies exposed to starvation on agar plates (n = 5).
